# Non-optimal codon usage is critical for protein structure and function of the master general amino acid control regulator CPC-1

**DOI:** 10.1101/2020.09.11.294470

**Authors:** Xueliang Lyu, Yi Liu

**Author notes:** Corresponding author: Yi Liu, Department of Physiology, ND13.214A, University of Texas Southwestern Medical Center, 5323 Harry Hines Blvd. Dallas, TX 75390-9040.

## Abstract

Under amino acid starvation condition, eukaryotic organisms activate a general amino acid control response. In *Neurospora crassa*, Cross Pathway Control-1 (CPC-1), the ortholog of the *Saccharomyces cerevisiae* bZIP transcription factor GCN4, functions as the master regulator of the general amino acid control response. Codon usage biases are a universal feature of eukaryotic genomes and are critical for regulation of gene expression. Although codon usage has also been implicated in the regulation of protein structure and function, genetic evidence supporting this conclusion is very limited. Here we show that *Neurospora cpc-1* has a non-optimal NNU-rich codon usage profile that contrasts with the strong NNC codon preference in the genome. Although substitution of the *cpc-1* NNU codons with synonymous NNC codons elevated CPC-1 expression in *Neurospora*, it altered CPC-1 degradation rate and abolished its amino acid starvation-induced protein stabilization. The codon-manipulated CPC-1 protein also exhibited different sensitivity to limited protease digestion. Furthermore, CPC-1 functions in rescuing the cell growth of the *cpc-1* deletion mutant and activating the expression of its target genes were impaired by the synonymous codon changes. Together, these results reveal the critical role of codon usage in regulating of CPC-1 expression and function, and establish a genetic example of the importance of codon usage in protein structure.

**Abstract importance:** General amino acid control response is critical for organisms to adapt to amino acid starvation condition. The preference to use certain synonymous codons are a universal feature of all genomes. Synonymous codon changes were previously thought to be silent mutations. In this study, we show that the *Neurospora cpc-1* gene has an unusual codon usage profile compared to other genes in the genome. We found that codon optimization of the *cpc-1* gene without changing its amino acid sequence resulted in elevated CPC-1 expression, altered protein degradation rate and impaired protein functions due to changes in protein structure. Together, these results reveal the critical role of synonymous codon usage in regulating of CPC-1 expression and function, and establish a genetic example of the importance of codon usage in protein structure.

## INTRODUCTION

Transcriptional regulation allows organisms to respond to changes in environmental conditions. Under amino acid starvation conditions, fungi activate a general amino acid control response that induces expression of genes involved in amino acid biosynthesis (1–4). The signal transduction pathways that mediate these responses are similar in eukaryotic cells from yeast to mammals. In the budding yeast *Saccharomyces cerevisiae* and the filamentous fungus *Neurospora crassa*, bZIP transcription factors GCN4 and Cross-Pathway Control-1 (CPC-1), respectively, are the master transcriptional regulators that activate amino acid biosynthetic genes in response to amino acid limiting conditions (1–6). Like GCN4, CPC-1 binds to the 5’-TGA(C/G)TCA-3’ motifs in target gene promoters to activate transcription (1–3, 5).

A mechanism involving upstream open reading frames (uORFs) in *GCN4* modulates GCN4 protein production (1, 2). Under normal conditions, the translation of the uORFs prevents translation initiation from the *GCN4* ORF, resulting in the suppression of GCN4 expression. Under amino acid starvation conditions, however, the scanning 40S ribosomes bypass the uORFs and initiate translation at the downstream *GCN4* ORF, resulting in the induction of GCN4 protein expression. This type of uORF-mediated mechanism is conserved in general amino acid control responses from fungi to mammals: Translational induction of CPC-1 in *Neurospora* and of ATF4 in mammals is controlled by this mechanism (7, 8). We recently showed that impaired tRNA I34 modification also triggers an amino acid starvation-like response (9). Ribosome profiling experiments, which were used to monitor ribosome occupancy on translating mRNAs, showed that there were many more ribosomes bypassing the two *cpc-1* uORFs and translating the downstream open reading frame region when tRNA I34 modification was suppressed (9).

Post-translational regulation of GCN4 stability is another mechanism that contributes to its up-regulation in response to amino acid starvation (1). GCN4 is very unstable when yeast cells are cultured in rich medium with a half-life of several minutes, but its degradation becomes much slower under amino acid starvation conditions (10, 11). The degradation of GCN4 is mediated by the proteasome ubiquitination pathway and is dependent on its phosphorylation by cyclin-dependent kinases (11–13). It is not clear whether amino acid availability also regulates CPC-1 stability in *Neurospora*.

Due to the degeneracy of genetic code, most amino acids are encoded by two to six synonymous codons. Codon usage bias, the preference for certain synonymous codons for almost all amino acids, is found in all genomes examined (14–17). Codon usage bias is an important determinant of gene expression levels in both eukaryotes and prokaryotes (18–21). We and other groups previously showed that codon usage regulates translation elongation speed: Common codons enhance the local rate of translation elongation, whereas rare codons slow down translation elongation (22–25). Rare codons preferentially cause ribosome stalling on an mRNA during translation, and this can result in premature translation termination and reduce translation efficiency (22, 24, 26). Furthermore, codon usage bias can regulate gene expression by affecting transcription (27–31).

In addition to the effect of codon usage on gene expression (32), accumulating biochemical and genetic evidence suggests that codon usage can also influence the co-translational protein folding process through its effects on translation elongation speed, which influences the time available for co-translational folding (19, 22, 26, 28, 33–46). In *Escherichia coli* (*E. coli*), it was previously shown that codon usage can affect protein activity and structures of some overexpressed proteins (37, 38, 40, 43, 47). In eukaryotes, a synonymous single-nucleotide polymorphism of the human *MDR1* gene was previously shown to cause altered protein activity of the MDR1 protein transiently overexpressed in human cells, suggesting the involvement of codon usage in eukaryotic protein folding (39). More recently, codon usage was also shown to influence protein activity and/or structures of several other human proteins (28, 45, 48–50). However, these previous studies relied on protein overexpression, which could also influence protein folding in cells, and the impacts of codon usage on protein structure/function were often modest.

By studying the circadian clock genes in *Neurospora* and *Drosophila*, we previously demonstrated that the codon usage of circadian clock gene *frq* in *Neurospora* and *Per* in *Drosophila* plays a major role in determining the protein structure and function *in vivo* (19, 36). Importantly, these studies did not use protein overexpression and the functional impacts of codon usage on protein function and structure were very robust in these genetic systems, thus confirming the physiological role of codon usage in protein folding in eukaryotic systems. Furthermore, genome-wide correlations between gene codon usage and predicted protein structures have been observed in prokaryotes and eukaryotes, suggesting that the codon usage functions as universal code to broadly modulate protein folding (33, 51–53). However, there are currently only very few genetic examples that demonstrate the robust physiological influence of codon usage on protein folding and function (19, 36, 38).

*N. crassa* has a strong codon usage bias for C/G at wobble positions (33, 54), but we observed that the *cpc-1* gene has an abnormal NNU-rich codon usage bias. Amino acid starvation triggers the stabilization of yeast GCN4, and we observed a similar starvation-induced stabilization of CPC-1 protein. By changing the NNU codons of *cpc-1* to synonymous NNC codons, we demonstrated that the codon usage of *cpc-1* is required for CPC-1 stabilization in response to amino acid starvation, and that it is critical for the CPC-1 structure and function *in vivo*. Together, our results demonstrate the role of codon usage in controlling CPC-1 expression and function, and establish another genetic example of the importance of codon usage in protein folding.

## RESULTS

### Abnormal codon usage profile of *cpc-1*

Examination of the *N. crassa cpc-1* gene revealed that it has an unusual codon usage profile. The *Neurospora* genome has a strong preference for NNC codons in every ADAT-related codon family (the A34 positions of their corresponding tRNAs can be converted to I34 by adenosine deaminases acting on tRNAs, known as ADATs) and for NNC/NNG codons in other codon families (33, 54). In contrast, for *cpc-1*, NNU codons are the most preferred codons for five of the eight ADAT-related codon families (Ala, Pro, Arg, Ser, and Val) (Figure 1A). For Leu codons of *cpc-1*, the normally preferred CUC codon is one of the least used codons; it has a lower usage frequency than CUU. The usage frequency of ACU, which codes for Thr, is also higher in *cpc-1* than the genome average (Figure 1A). Interestingly, the genome preferred NNC codons are also not the preferred codons for the majority of the ADAT-related codon families in homologous *cpc-1* genes in *Neurospora tetrasperma, Sordaria macrospora* and *Aspergillus nidulans* (Supplementary Figure S1), suggesting that non-optimal *cpc-1* codon usage is conserved. These results raised the possibility that the non-optimal nature of the *cpc-1* codon usage profile is functionally important.

**Figure 1.**
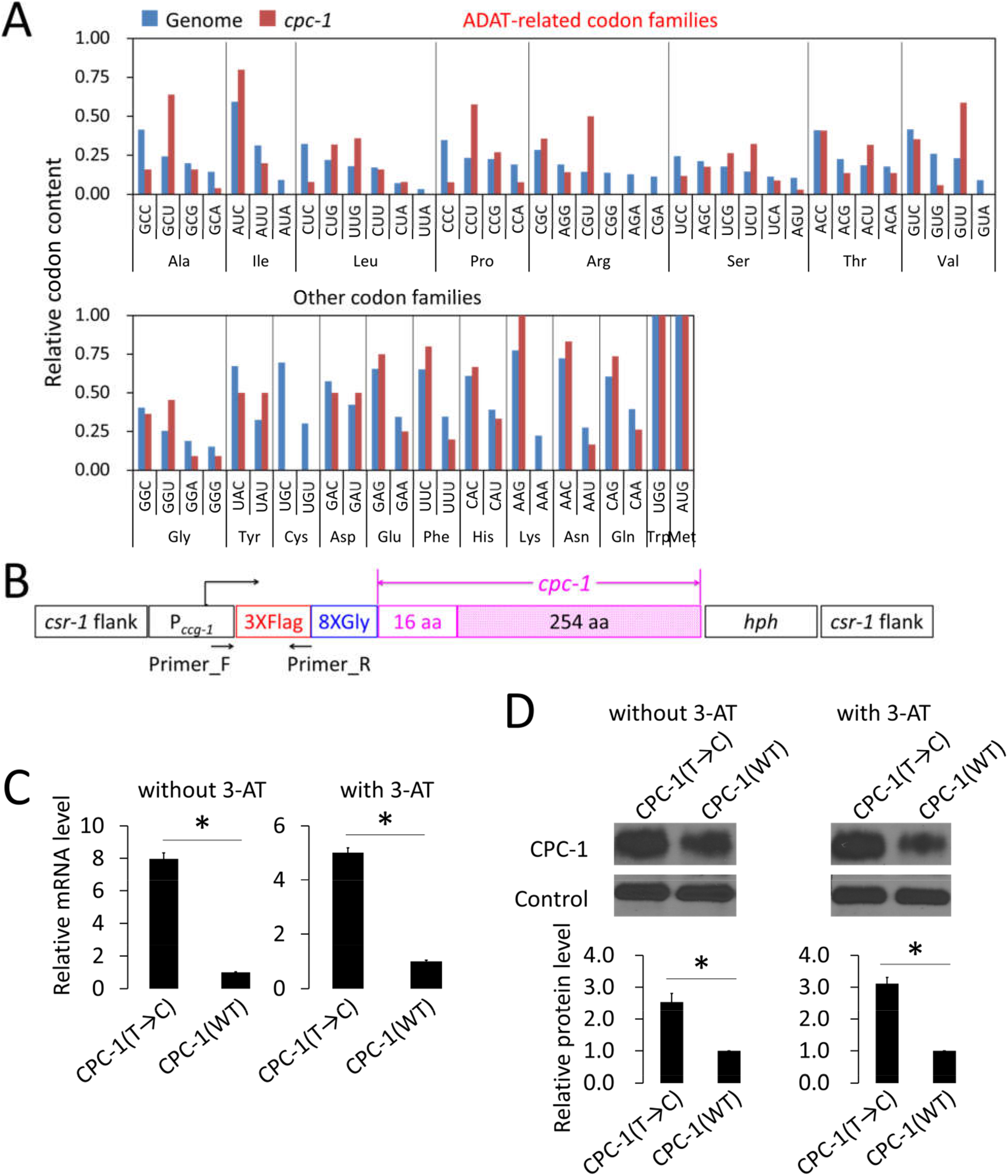
The ADAT-related NNU-rich codon usage of *cpc-1* contributes to the regulation of CPC-1 production. **(A)** The codon usage of *cpc-1* represented by the relative codon contents in each codon family compared to the genome-wide average codon usage. **(B)** Graphical representation of the constructs used for the expression of *cpc-1*(WT) and optimized *cpc-1*(T→C). To avoid the influences of the *cpc-1* uORFs and 5’ UTR, the *ccg-1* promoter (P_*ccg-1*_) and its 5’ UTR were used to drive the expression of *cpc-1*. The constructs were integrated into the *csr-1* locus in the *N. crassa* genome by homologous recombination. To avoid the influence of codon optimization on translation initiation, the first 50 codons (including the codons of 3×Flag, 8×Glycine, and 16 codons at the N terminus of *cpc-1*) were kept the same in both constructs. In *cpc-1*(T→C), 71 of the 270 ADAT-related NNU codons were changed to the most preferred NNC codons of *N. crassa* genome. For details see Supplementary Figure S2. **(C)**The relative mRNA levels of *cpc-1*(WT) and *cpc-1*(T→C) detected by qRT-PCR in the host strain cultured in 2% glucose medium with or without 5 mM 3-aminotriazole (3-AT). The *cpc-1* mRNA levels were normalized to that of *β-tubulin* gene (NCU04054). Primers used for qRT-PCR were designed to the 5’ region of the transcript, which is common to both of the constructs (as shown in B) to ensure the same amplification efficiency. The *cpc-1*(WT) transcript level was set as 1.0. **(D)** Upper panel: Western blot analysis of CPC-1 expressed in the *cpc-1*(WT) and *cpc-1*(T→C) strains cultured in 2% glucose medium with or without 5 mM 3-AT. A non-specific constitutive band detected by the anti-FLAG antibody was used as control. Lower panel: Densitometric analyses of the CPC-1 levels from three independent experiments. The CPC-1 protein level produced from *cpc-1*(WT) was set as 1.0. Data in panels C and D are means ± SD (n = 3). *, P < 0.05, as determined by Student’s two-tailed t-test.

### Regulation of CPC-1 expression by *cpc-1* codon usage

The unusual NNU-rich codon usage profile of *cpc-1* suggests that it may play a biological role in regulating CPC-1 expression or function. To test this hypothesis, we created two versions of the *cpc-1* ORF: *cpc-1*(WT), in which all native codons were maintained; *cpc-1*(T→C), in which all eight ADAT-related NNU codons (except for the codons for the N-terminal 16 amino acids) were substituted synonymously with the genome-preferred NNC codons without altering the amino acid sequence (Supplementary Figure S2). Thus, the *cpc-1*(T→C) has more optimal codons than the wild-type (WT) *cpc-1*. We expressed 5’ epitope-tagged versions of *cpc-1*(WT) and *cpc-1*(T→C) ORFs under the control of the *ccg-1* promoter and the *ccg-1* 5’ untranslated region (5’ UTR) to exclude the effect of the uORFs and the 5’ UTR of *cpc-1* on translation. To minimize the potential impact of codon usage on translation initiation, the N-terminal regions (which include 3×Flag, an 8×Gly linker, and the codons for the initial 16 N-terminal amino acids of CPC-1) of the two versions of *cpc-1* were identical (Figure 1B). Constructs containing the *cpc-1* transgenes were individually transformed into the *Neurospora* strain 87-3 at the targeted *csr-1* locus. Homokaryotic transformant strains were cultured in 2% glucose medium with or without 3-aminotriazole (3-AT). 3-AT treatment results in amino acid starvation in *Neurospora* because it is a competitive inhibitor of the product of *his-3* gene, which is an enzyme required for histidine biosynthesis (6, 55–57). Because of the cross-pathway control in *Neurospora*, depletion of one amino acid leads to a general amino acid starvation response (3).

Gene codon optimization usually results in increased mRNA and protein levels in *Neurospora* (27, 58, 59). As expected, the mRNA levels of *cpc-1*(T→C) were significantly higher than that of the *cpc-1*(WT) when they were expressed in the host strain cultured in 2% glucose medium with or without 3-AT (Figure 1C). Similarly, the CPC-1 protein levels were also up-regulated in the *cpc-1*(T→C) strain (Figure 1D). These results suggest that the NNU-rich codon usage profile of *cpc-1* suppresses CPC-1 expression.

### Codon usage of *cpc-1* and culture conditions affect CPC-1 protein stability

GCN4, the ortholog of CPC-1 in *S. cerevisiae*, is rapidly degraded under rich nutrient conditions but it is stabilized under amino acid starvation conditions, a response that contributes to GCN4 up-regulation after amino acid starvation (1, 10, 11, 60). To determine whether CPC-1 protein stability is affected by amino acid starvation and codon usage, we compared CPC-1 turnover rates after the addition of the protein synthesis inhibitor cycloheximide (CHX) in the *cpc-1*(WT) and *cpc-1*(T→C) strains grown in 2% glucose medium with and without 3-AT. 3-AT treatment resulted in marked stabilization of CPC-1 in the *cpc-1*(WT) strain (Figure 2), suggesting that as with GCN4 in yeast (1), protein stability is altered under amino acid starvation conditions. In the *cpc-1*(T→C) strain, however, CPC-1 was more stable than that in the *cpc-1*(WT) strain when the strains were grown in 2% glucose medium without 3-AT. Furthermore, the stabilization of CPC-1 observed in the *cpc-1*(WT) strain upon 3-AT treatment was not observed in the *cpc-1*(T→C) strain (Figure 2). These results indicate that the NNU-biased *cpc-1* codon usage plays an important role in regulating CPC-1 protein stability under amino acid starvation conditions.

**Figure 2.**
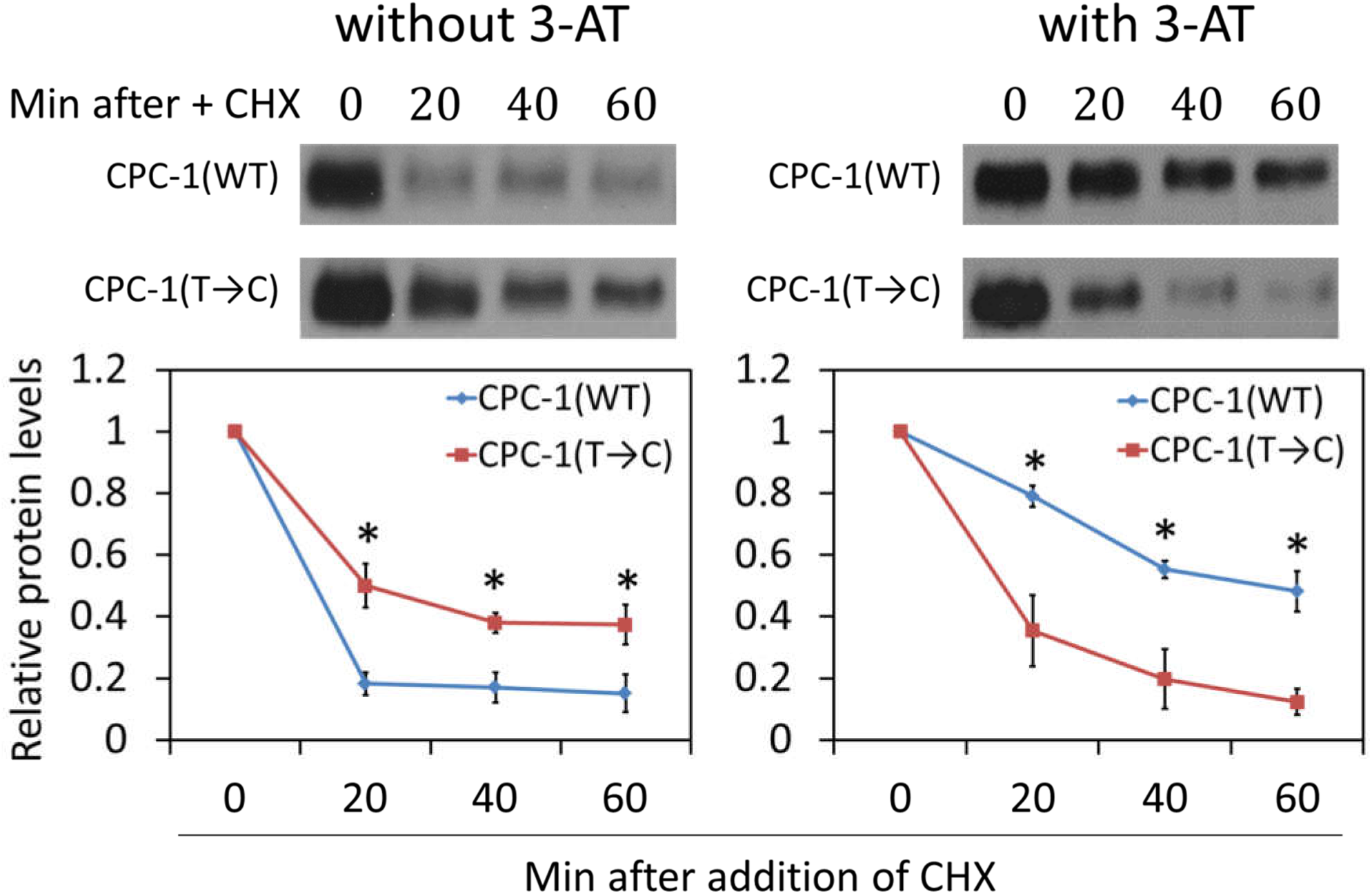
Codon usage optimization alters CPC-1 stability in response to amino acid starvation. Upper panels: Representative western blots showing the CPC-1 protein levels in *cpc-1*(WT) and *cpc-1*(T→C) strains grown in 2% glucose medium with or without 5 mM 3-AT. Cycloheximide (CHX, 10 μg ml^-1^) was added at time 0 and cultures were harvested at the indicated time points. Lower panels: Densitometric analyses of the western blot experiments described in the upper panels. Data are means ± SD (n = 3). *, P < 0.05, as determined by Student’s two-tailed t-test.

### *cpc-1* codon usage affects CPC-1 structure and function

The effect of *cpc-1* codon usage on CPC-1 protein stability raised the possibility that proper co-translational folding of CPC-1 depends on codon usage as observed for other *Neurospora* proteins (19, 22, 24). To examine this possibility, we performed a limited trypsin digestion assay to probe the structure differences of CPC-1 proteins in the *cpc-1*(WT) and *cpc-1*(T→C) strains. The freshly isolated protein extracts of the *cpc-1*(WT) and *cpc-1*(T→C) strains were treated with trypsin, and the levels of full-length CPC-1 were determined by western blot analyses as a function of digestion time. When the cultures were grown in 2% glucose medium without 3-AT, CPC-1 isolated from the *cpc-1*(T→C) strain was significantly more resistant to trypsin digestion than that isolated from the *cpc-1*(WT) strain, but it was more sensitive to trypsin digestion after 3-AT treatment (Figure 3). These results indicate that codon usage influences CPC-1 protein structure.

**Figure 3.**
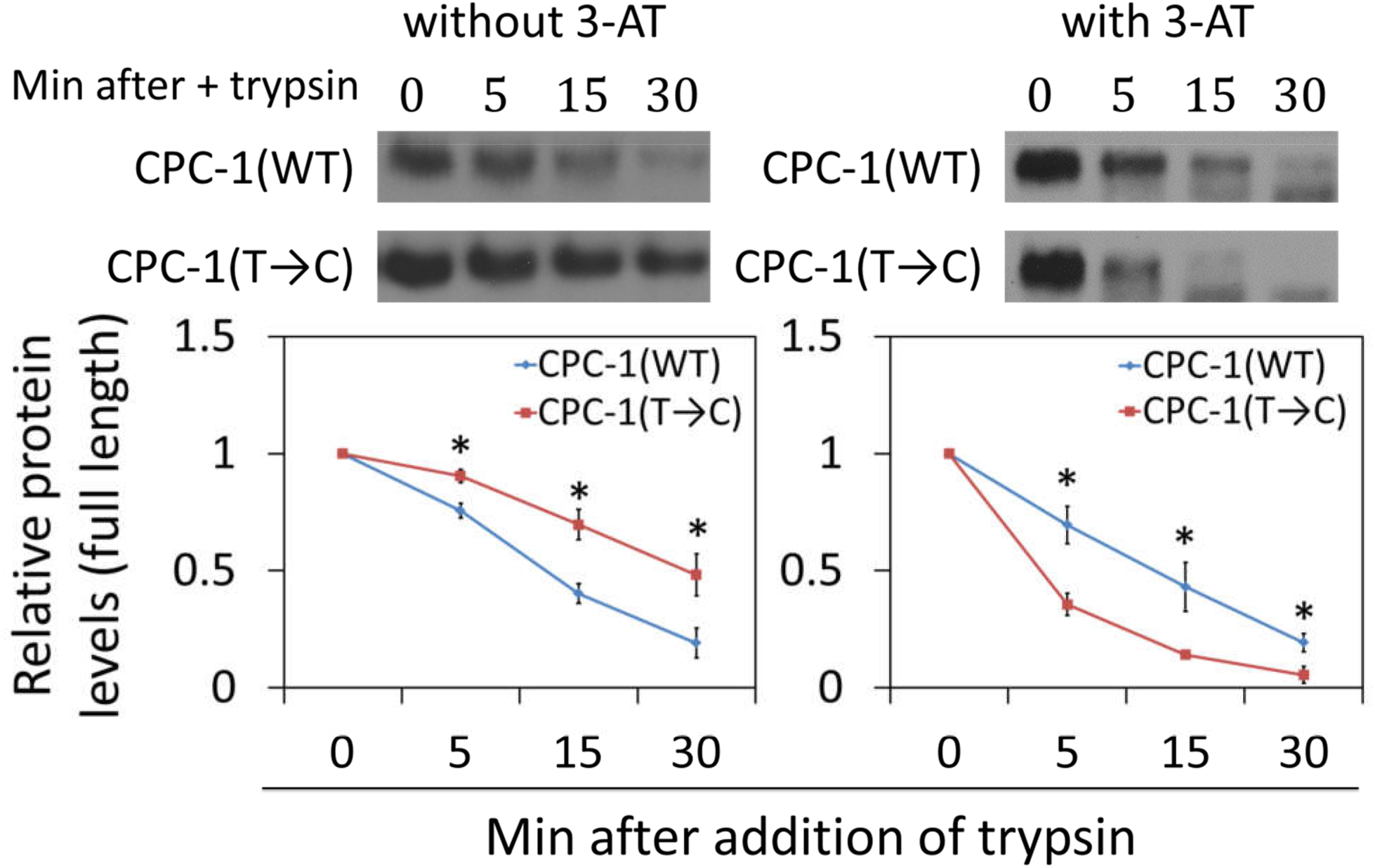
Codon usage affects CPC-1 co-translational folding. **(Top)** Western blots of CPC-1 expression in the *cpc-1*(WT) and *cpc-1*(T→C) strains cultured in 2% glucose medium in the absence or presence of 5 mM 3-AT. Trypsin (0.25 μg/ml) was added into the freshly isolated protein extracts, and protein samples were analyzed at the indicated time points. **(Bottom)** Densitometric analyses of the full-length CPC-1 levels from the experiments above. Data are means ± SD (n = 3). *, P < 0.05, as determined by Student’s two-tailed t-test.

It is important to note that the *ccg-1* promoter-driven *cpc-1* expression did not result in its overexpression. In fact, we found that the *cpc-1* mRNA level under the control of *ccg-1* promoter and 5’UTR in a *cpc-1* knock-out strain (*cpc-1*Δ) was actually much lower than that of the endogenous *cpc-1* level in a WT strain (supplementary Figure S3). Thus, the effect of codon usage on CPC-1 structure is not due to its overexpression.

To determine whether the structural differences caused by codon usage result in changes in protein function, we introduced the *cpc-1*(WT) and *cpc-1*(T→C) constructs individually into the *cpc-1*Δ strain. We then compared the abilities of these two constructs to rescue the growth defect of the *cpc-1*Δ mutant under amino acid starvation conditions. Under normal growth conditions, the WT and *cpc-1*Δ strains had similar growth rates, but the *cpc-1*Δ strains expressing the *cpc-1*(WT) or *cpc-1*(T→C) had a slightly but significantly slower growth rate in constant light at room temperature (Figure 4A). In constant light, *cpc-1*(WT) and *cpc-1*(T→C) are constitutively expressed, and *cpc-1* translation is not regulated by the uORFs due to the use of the *ccg-1* 5’ UTR in the transgene strains. The reduced growth rate in these strains is consistent with the known role of GCN4 as a repressor of protein synthesis (61). In the presence of 3-AT, however, the growth rate of *cpc-1*Δ strain was dramatically reduced, compared to the WT strain in a race tube assay (Figure 4B). The growth phenotype was drastically improved in the *cpc-1*(WT) strain, indicating a functional rescue of the *cpc-1*Δ strain by the *cpc-1*(WT) transgene. The growth rate of the *cpc-1Δ, cpc-1*(T→C) strain, however, was much slower than that of the *cpc-1*Δ, *cpc-1*(WT) strain in the presence of 3-AT (Figure 4B), indicating that the T→C codon usage profile changes impaired the CPC-1 protein function even though it increased CPC-1 protein level (Figure 1D).

**Figure 4.**
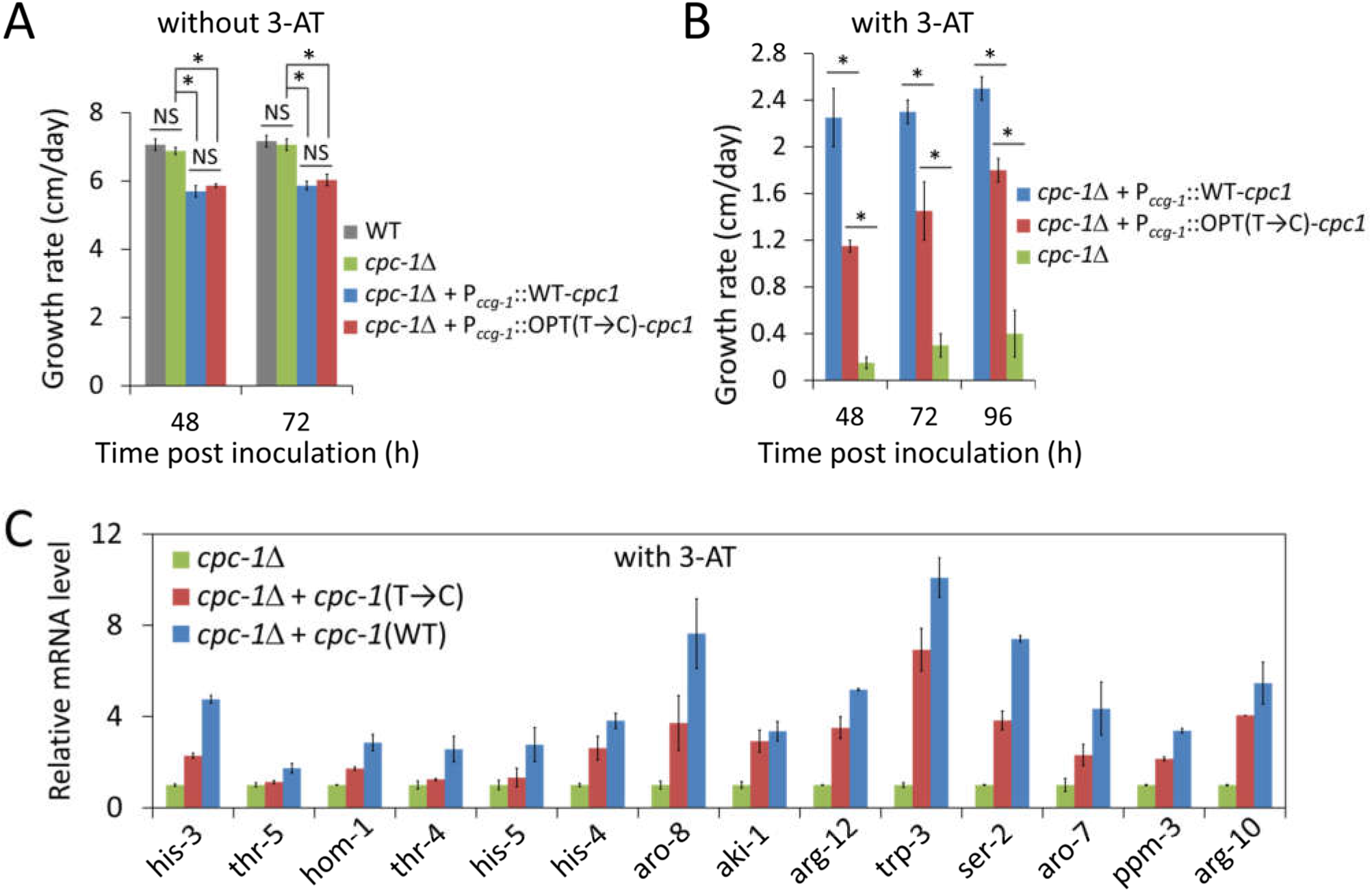
Optimization of *cpc-1* codon usage impairs CPC-1 biological functions. **(A)** Growth rates of the WT strain, the *cpc-1*Δ strain, and the *cpc-1*Δ strains expressing *cpc-1*(WT) or *cpc-1*(C→T) after 48 and 72 h as determined in race tube assays without 3-AT. *, P < 0.05; NS, not significant; as determined by Student’s two-tailed t-test. **(B)** Growth rates of the *cpc-1*Δ strain and the *cpc-1*Δ strains expressing *cpc-1*(WT) or *cpc-1*(C→T) after 48, 72, and 96 h as determined in race tube assays with 5 mM 3-AT. **(C)** The relative mRNA levels of selected CPC-1 target genes in the indicated strains. The relative mRNA level of each gene was determined by qRT-PCR, and their expression levels were normalized to that of *β-tubulin* gene (NCU04054). The mRNA level of each gene in the *cpc-1*Δ strain was set as 1.0. Data in panels A, C, and D are means ± SD (n = 3).

To further confirm this conclusion, we compared the mRNA levels of 14 genes regulated by CPC-1 that are involved in amino acid biosynthesis (3, 9). The mRNA levels of these CPC-1 target genes in the *cpc-1*Δ, *cpc-1*Δ, *cpc-1*(WT) and *cpc-1*Δ, *cpc-1*(T→C) strains treated with 3-AT were determined. The mRNA levels of all the 14 genes were dramatically up-regulated in the *cpc-1Δ, cpc-1*(WT) strain compared to those in the *cpc-1*Δ strain (Figure 4C). As expected, the mRNA induction of these genes was reduced in the *cpc-1*Δ, *cpc-1* (T→C) strain (Figure 4C). Together, these results demonstrate that the NNU-biased codon usage profile of *cpc-1* plays an important role in determining CPC-1 protein structure and function *in vivo*.

## DISCUSSION

CPC-1 is the master transcription regulator of gene expression in *Neurospora* in response to amino acid starvation (3, 7, 55, 62). Like the situation of its yeast ortholog GCN4, the expression of CPC-1 is also translationally regulated by a mechanism involving uORFs (7, 9). The translation of *cpc-1* can be translationally activated by bypassing its uORFs under amino acid starvation condition. Like GCN4 (10, 11), here we showed that 3-AT treatment triggers CPC-1 stabilization, which also contributes to CPC-1 accumulation under amino acid starvation. Although the mechanism of CPC-1 degradation is not known, it is possible that like GCN4, amino acid starvation regulates posttranslational modification of CPC-1, which affects its degradation by the ubiquitin-proteasome pathway (11–13).

In this study, we demonstrated that the non-optimal codon usage profile of *cpc-1* has a major impact on the structure and function of CPC-1. By changing the *cpc-1* codon usage from the NNU-rich profile to the NNC-rich profile typical of the *Neurospora* genome, we showed that codon usage is critical for CPC-1 protein structure and function. This conclusion was supported by several lines of evidence. First, the codon manipulation altered CPC-1 protein degradation rate and abolished amino acid starvation induced CPC-1 stabilization (Figure 2), suggesting that codon usage can affect CPC-1 structure. Second, the codon optimization altered CPC-1 sensitivity to limited trypsin digestion, indicating that codon optimization affected protein structure (Figure 3). Third, in the presence of 3-AT, the CPC-1(T→C) was less stable and more sensitive to trypsin digestion than the CPC-1 (WT) (Figure 2 & 3), suggesting that codon usage-mediated structure changes of CPC-1 affected its ability to be regulated by potential posttranslational mechanisms triggered by amino acid starvation conditions. Fourth, despite the up-regulation of CPC-1 protein levels in *Neurospora* upon codon optimization, the expression of *cpc-1*(T→C) did not as effectively rescue the growth defects of the *cpc-1*Δ strain under amino acid starvation condition as that of *cpc-1*(WT) did. Finally, the impaired CPC-1 function of the *cpc-1*(T→C) strain was further indicated by the reduced induction of CPC-1 target genes in response to amino acid starvation. It is important to note that the codon optimized *cpc-1* did not cause its overexpression (supplementary Figure S3). Thus, our study established a critical physiological role of codon usage in regulating CPC-1 structure and function.

In addition, our analyses established another *in vivo* example of the influence of codon usage on protein structure. Due to the role of codon usage in regulating translation elongation rate, codon usage was proposed to influence the co-translational protein folding process (19, 22, 26, 28, 33–38). However, genetic evidence in support of such a role of codon usage is quite limited. By studying the codon usage function of the circadian clock genes *frequency* in *Neurospora* and *Period* in *Drosophila*, we previously showed that codon usage plays an important role in affecting the structures, and therefore functions, of these two proteins *in vivo* (19, 36). Similar to *frequency* and *Period* genes, *cpc-1* is enriched in non-optimal codons. Additionally, as with FRQ and PER proteins, most regions of the CPC-1 protein are predicted to be intrinsically disordered. Our findings are consistent with the hypothesis that the co-translational protein folding process is sensitive to codon usage-mediated translation elongation kinetics and that this process is regulated to ensure proper functioning of the proteins with intrinsically disordered domains. Further supporting this, we and others previously showed that non-optimal codon usage correlated with predicted unstructured domains genome-widely in *Neurospora* and other organisms (33, 52). The structure of the DNA binding domain of yeast GCN4, the CPC-1 ortholog, was previously shown to be flexible (63–65). GCN4 exhibits a concentration-dependent α-helical transition: The transition of the GCN4 basic region from an unfolded to a folded conformation depends on its accessibility to DNA binding site (65). Such properties may make it more sensitive to the co-translational folding process.

Taken together, our results suggest that the unusual codon profile of *cpc-1* represents another example of evolutionary adaption that results in its optimal protein structure and function in response to environmental changes.

## MATERIALS and METHODS

### Strains and growth conditions

*N. crassa* 87-3 strain (*bd, a*) was used as the control and further used as the host strain for the expression of various versions of *cpc-1* unless otherwise specified. For the growth rate assay, the FGSC 4200 (*a*, WT) strain was used as the control. The *cpc-1*Δ strain was obtained from the *Neurospora* knock-out library (66). Liquid cultures were grown in 2% glucose medium (1 × Vogel’s, 2% glucose) or 0.1% glucose medium (1 × Vogel’s, 0.1% glucose and 0.17% arginine). Race tube medium contained 1 × Vogel’s, 0.1% glucose, 0.17% arginine, 50 ng ml^-1^ biotin, and 1.5% agar. All the strains were cultured on slants containing 1 × Vogel’s, 2% sucrose, and 1.5% agar before performing various experiments. All the strains were cultured under constant light at room temperature.

### Plasmid constructs

For gene expression at the *csr-1* locus in *N. crassa*, a hygromycin B resistance gene (*hph*) was inserted downstream of the *ccg-1* promoter of a parental plasmid Pcsr1 to create a new plasmid Pcsr1-hyg. Pcsr1-hyg is a *csr-1*-targeting expression vector with an expression cassette in which P_ccg−1_ and *hph* flank the gene of interest, and this cassette is flanked by two *csr-1*-related fragments that serve as the double recombination sites (67). When this plasmid is transformed into *N. crassa* cells, it is integrated into the *csr-1* gene locus by replacing *csr-1* with the expression cassette by double homologous recombination. The resulting transformants are screened for both hygromycin B (200 μg ml^−1^) and cyclosporin A (5 μg ml^−1^) resistance conferred by the presence of *hph* and the absence of *csr-1*, respectively. The efficiency and accuracy of this approach were very high (> 90% positive transformants). In this study, two versions of *cpc-1*(WT) and *cpc-1*(T→C) with a 3×Flag tag and an 8×Glycine linker at the N-termini were introduced into the Pcsr1-hyg construct. The resulting constructs were transformed into host strains by electroporation. Homokaryon strains were obtained by microconidia purification.

### Protein stability and limited trypsin digestion assays

For protein stability assay, cycloheximide (CHX) working concentration and experimental procedures were the same as previously described (19). For culture conditions, fresh conidia (one week post inoculation on slants) of the host strains were cultured in 50 ml 2% glucose medium in plates at room temperature for two days. The cultures were cut into small discs with a diameter of 1 cm, and then the discs were transferred into flasks with the same liquid medium and were grown with orbital shaking (200 rpm) for one more day before adding CHX (final concentration 10 μg ml^-1^). For the sample treated with 3-AT, the culture discs in 2% glucose medium were treated with 5 mM 3-AT for 8 h before sample collection. Cells were collected at the indicated time points after adding CHX. For the limited trypsin digestion assay, the culture conditions and sample collection procedures were the same as the above except for the addition of CHX. The working concentration of trypsin was 0.25 μg/ml. Protein extraction and western blot analyses were performed as previously described (68). Equal amounts of total proteins (100 μg) were loaded into each lane of 7.5% SDS-PAGE gels containing 37.5:1 acrylamide/bisacrylamide. The primary and secondary antibodies used for detecting the 3×Flag were monoclonal ANTI-FLAG® M2 antibody produced in mouse (Sigma-Aldrich, Cat NO.: F3165) and Goat Anti-Mouse IgG (H + L)-HRP Conjugate (Bio-Rad, Cat NO.: 170-6516), respectively. Densitometry was performed using Image J.

### Quantitative reverse transcription PCR (qRT-PCR) and mRNA-seq

For qRT-PCR, the sample collection procedures are the same as described in the protein stability assay section except that CHX was not added. For cultures treated with 3-AT as indicated in figures, the liquid cultures were treated with 5 mM 3-AT for 8 h before sample collection. RNA extraction and qRT-PCR were performed as previously described (69). *β-tubulin* (NCU04054) was quantified as an internal control. Primers used for qRT-PCR are listed in Supplementary Table S1. The relative mRNA levels of *cpc-1* in WT, *cpc-1*Δ and *cpc-1*Δ, *cpc-1*(WT) strains under amino acid starvation condition were measured by their RPKM (Reads Per Kilobase Per Million) values from our high-throughput mRNA sequencing data. The mRNA-seq libraries in this study were generated from cultures in 2% glucose medium with 5 mM 3-AT treatment for 8 h before sample collection. The sample collection procedures are the same as described in the protein stability assay section except that CHX was not added. Total RNAs were extracted using Trizol reagents (Invitrogen) and treated with DNase (Turbo DNase, Ambion). The libraries were prepared according to the NEBNext Ultra Kits for RNA and sequenced by Illumina HiSeq2000. mRNA-seq experiments were performed by Joint Genome Institute (JGI) on Illumina novaseq platform. The raw and processed sequencing data have been submitted to the NCBI Gene Expression Omnibus under accession number GSE150287.

### Codon manipulation and data collection from databases

The codons of *cpc-1* were optimized based on the *N. crassa* codon usage frequency from the Codon Usage Database (https://www.kazusa.or.jp/codon/cgi-bin/showcodon.cgi?species=5141). The mutated sites for the optimized *cpc-1*(T→C) are shown in Supplementary Figure S2.

## ACKNOWLEDGMENTS

We thank the members of our laboratory for assistance. This work is supported by grants from the National Institutes of Health (R35GM118118), and the Welch Foundation (I-1560) to Yi Liu. Xueliang Lyu is partially supported by National Natural Science Foundation of China (31701735) and the International Postdoctoral Exchange Fellowship Program 2017 by the Office of China Postdoctoral Council ([2017]32).

## SUPPLEMENTARY FIGURES

**Figure S1. The codon usage profiles for ADAT-related codon families of *cpc-1* homologous genes in *Neurospora tetrasperma, Sordaria macrospora* and *Aspergillus nidulans***. The relative codon contents compared to the genome-wide average codon usage in each codon family are shown.

**Figure S2. Sequence alignment of the coding sequences of *cpc-1*(WT) and optimized *cpc-1*(T→C). * indicate conserved sites.**

**Figure S3. The relative *cpc-1* mRNA levels detected in the indicated strains.** The relative expression levels of *cpc-1* were measured by RPKM (Reads Per Kilobase Per Million) values. Only reads mapped to the CDS region of *cpc-1* were taken into account.

**Table S1: Sequences of primers used in this study.**

